# Social effects of rabies infection in male vampire bats (*Desmodus rotundus*)

**DOI:** 10.1101/2022.06.06.495012

**Authors:** Elsa M. Cárdenas Canales, Sebastian Stockmaier, Eleanor Cronin, Tonie Rocke, Jorge E. Osorio, Gerald G. Carter

**Author notes:** **Corresponding authors:** EMCC, SS. co-first authors (Elsa M. Cárdenas Canales, Sebastian Stockmaier).

## Abstract

Rabies virus (RABV) transmitted by the common vampire bat (*Desmodus rotundus*) poses a threat to agricultural development and public health throughout the Neotropics. The ecology and evolution of rabies host-pathogen dynamics are influenced by two infection-induced behavioral changes. RABV-infected hosts often exhibit increased aggression which facilitates transmission, and rabies also leads to reduced activity and paralysis prior to death. Although several studies document rabies-induced behavioral changes in rodents and other dead-end hosts, surprisingly few studies have measured these changes in vampire bats, the key natural reservoir throughout Latin America. Here, we take advantage of an experiment designed to test the safety and efficacy of an oral rabies vaccine in captive male vampire bats to quantify for the first time how rabies affects allogrooming and aggressive behaviors in the vampire bat. Compared to non-rabid vampire bats, rabid individuals reduced their allogrooming prior to death, but we did not detect increases in aggression among bats. To put our results in context, we review what is known and what remains unclear about behavioral changes of rabid vampire bats.

## Introduction

Rabies virus (RABV) transmitted by the blood-feeding common vampire bat (*Desmodus rotundus*) creates a substantial burden for agricultural development and public health throughout Latin America, with deadly rabies outbreaks occurring in livestock [1–3] and humans [4,5]. RABV is transmitted by direct contact between the virus-laden saliva of the infected bat and the other animal’s broken skin, eyes, or mucous membranes. RABV transmission can occur both among vampire bats and when they bite livestock, wildlife, or less frequently, humans, leading to cross-species transmission [2,6]. Recent studies combining mathematical modeling, RABV phylodynamics, and the ecology, demography, and dispersal of vampire bats have shown great potential to predict and mitigate these pathogen spillover events [6–9].

Surprisingly few studies, however, have directly investigated how RABV infection affects vampire bat behavior. In mustelids [10,11], canines [12,13], rodents [14], and humans [15], RABV can lead to paralysis without obvious increases in aggression before death (“paralytic” rabies), but it can also induce aggression and biting (“furious” rabies) which is likely to increase transmission to other hosts (pathogen manipulation) [16]. Given that aggressive interactions are commonly observed in vampire bats, especially among males [17–19], and that RABV is detectable in the saliva at the end of infection [20,21], increases in aggression in rabid vampire bats should enhance transmission. In a previous study [21], seven confirmed naturally RABV-exposed vampire bats showed no obvious symptoms and survived, while seven others presented two distinct disease outcomes. Three bats showed *furious* rabies presentation with hypersalivation, excess vocalizations, teeth chattering, aggression towards handlers and other bats, and irritability to light and sound. Four bats showed *paralytic* rabies presentation with social isolation, lethargy, and apparent respiratory distress. Although these anecdotal observations demonstrate both presentations are possible, they appear in some studies but not others (Table 1), and the relative probability of paralytic versus furious symptoms in rabid vampire bats remains unclear.

**Table 1:** Studies anecdotally describing rabies-induced changes in social behavior of vampire bats after injection or natural exposure.

| *Reference* | *Result* | *Method* | *Type of virus used* | *Comments* |
| --- | --- | --- | --- | --- |
| Present study | No increase in aggressive behavior observed, reduced social grooming | Behavioral sampling | Coyote variant | 40 male vampire bats. Some previously vaccinated. 15 bats rabies confirmed. |
| [25] | No aggression, grooming unobserved | Anecdotal observation | T-9/95 vampire bat field isolate from before 2001 (no detailed information). Study published in 2009. Isolate location unknown. | 10 vampire bats (7 males, 3 females). 4 bats confirmed positive. Signs of paralytic rabies in all 4. |
| [31] | No aggression, grooming unobserved | Anecdotal observation | CASS-88 vampire bat origin, isolated from infected cow brain in 1988. Study published in 1998. | 24 vampire bats (no information on sex). Some bats previously vaccinated with oral vaccine. 16 confirmed positive. Signs of paralytic rabies in 10/16. Others showed no obvious clinical signs. |
| [20] | No aggression, grooming unobserved | Anecdotal observation | CASS-88 vampire bat origin, isolated from infected cow brain in 1988. Study published in 2005. | 14 bats (6 males, 8 females). 11 died. Paralysis of wings and hind-legs prior to death in 3/11. Others showed depression, hypoactivity and anorexia. |
| [32] | No aggression, grooming unobserved | Anecdotal observation | CASS-88 vampire bat origin, isolated from infected cow brain in 1988. Study published in 2002. | Test of rabies vaccine. 9/10 control bats (vaccinated with saline) died of rabies (no information on sex). Altered reflexes, tremor, and paralysis were observed 72–24 hours before death in rabid bats. |
| [33] | No aggression, grooming unobserved | Anecdotal observation | Brldr2918 vampire bat field isolate from 1997. Study published in 2005. Study bat location and location of virus isolate are <100 km apart. | 10 bats died of RABV (no detailed information on sex/deaths). 8 showed signs of paralytic rabies, 2 showed no clinical signs |
| [34] | No aggression, grooming unobserved | Anecdotal observation | Brldr2918 vampire bat field isolate from 1997. Study published in 2008. No information on bat capture location | 10 bats died of RABV (no detailed information on sex/deaths) some were previously vaccinated (orally, fur-transmissible). All showed signs of paralytic rabies. |
| [21] | Aggression, grooming unobserved | Anecdotal observation | Natural exposure | Total of 14 confirmed rabid male bats. 7 showed no clinical signs, 3 the furious form and 4 the paralytic form. |
| [35] | Potential aggression, grooming unobserved | Anecdotal observation | Natural exposure | Introduction of wild bats to an existing captive colony (no information on sex). After 2 months, fighting started. Several bats from original colony were mutilated and tested for rabies. Suggested aggression of introduced bats. |
| [36] | Aggression, grooming unobserved | Anecdotal observation | Natural exposure | Aggression observed in naturally infected rabid bats. 14 of 24 bats observed showed clinical signs including hyperexcitability, aggressiveness, and paralysis before death. |

Besides biting, another possible transmission pathway is allogrooming, i.e., the licking and chewing of a conspecific’s fur and skin [22]. Allogrooming takes up about 3-5% of a bat’s active time [23], is sometimes targeted to wounds on the skin, and can reopen minor wounds (GGC, personal observation) creating potential for transfer of saliva. Allogrooming of the face and mouth is sometimes followed by regurgitations of ingested blood (e.g., [24]), which could also lead to RABV transmission [25]. No study has yet quantified changes in allogrooming in rabid bats.

During a study to evaluate a recombinant rabies vaccine candidate for vampire bats, we opportunistically measured rates of aggression and allogrooming in 40 captive male vampire bats that were experimentally infected with RABV. We then compared aggression and allogrooming in non-rabid bats to bats confirmed positive for rabies at death or the end of the study.

## Material and Methods

### Capture and care

We collected behavior data from 40 male common vampire bats that were part of a larger sample of bats used to test a viral-vectored recombinant mosaic glycoprotein rabies vaccine candidate. The bats were captured in the State of San Luis Potosí, México, July-Aug 2018, and transported to the U.S. Geological Survey National Wildlife Health Center in Madison, Wisconsin, USA (for details see [21]). Bats were individually marked by combinations of 0-4 bat bands (Porzana Limited, Icklesham, UK) on the right or left forearm.

### Experimental procedure

Bats were caged according to three treatments: (1) oral vaccination, (2) topical vaccination, or (3) placebo control. Treated bats remained caged together for ∼120 days before being challenged with RABV. One week prior to the challenge, we reassigned the bats into new groups, so individuals that received different treatments would be included in each cage and given time to acclimate (3 cages with 13, 13, and 14 bats each). All bats were challenged with a heterologous RABV variant (of coyote origin) at a dose of 10^3.3^ tissue culture infective dose (TCID_50_/mL), injected intramuscularly into each masseter muscle (50 µL on each side) in April 2019 (127 days post-vaccination). We began quantifying behaviors one day after the challenge. To confirm death by rabies, we performed a direct fluorescent antibody test for RABV in brain impression smears of bats following standard procedures [26]. To detect RABV shedding in the saliva, we collected oral swabs periodically from all individuals, daily if clinical signs were observed, and upon death. Swabs were tested using real-time PCR as described elsewhere [27,28].

### Behavioral data collection

After the bats were challenged with RABV, they were recorded using an infrared surveillance system (Amcrest 960 H/+) with a different camera pointed into each cage through a clear acrylic viewing window. In each of the three cages, we sampled behaviors three hours per night (at hours 0100, 0300, and 0500) during the most active period [29]. At every new-minute mark, an observer that was blind to the infection status of the bats stopped the video and recorded the presence or absence of either allogrooming or aggression within a 5-second time window and the identities of the actor and receiver (based on their unique combination of forearm bands). Allogrooming involves licking or chewing another bat’s fur or skin and often occurs with two bats allogrooming each other simultaneously (video S1). Aggressive events included biting and fighting (video S2), and a behavior we call “clinging” where a bat bites into another bat’s neck and clings onto it for a prolonged time period while the target is actively trying to shake off the aggressor (video S3).

### Statistical analysis

For each night we collected 180 presence/absence samples per group except for two nights when 53 samples and 29 samples were missing due to camera outages (resulting in a total of 18,818 behavioral samples). We counted the number of observed allogrooming and aggression events for each bat and divided it by the three sampled hours to estimate behavioral rates per hour. We estimated the 95% confidence intervals (CIs) around the mean rate for each of the two treatment groups, rabid and non-rabid, using bootstrapping (percentile method, 5,000 iterations, boot R package, [30]).

Because the exact timeline of infection was unclear, we plotted for every rabid bat an effect size (standardized mean difference between observed and expected) during increasingly large (nested) time periods, from 1 day to 15 days before death (excluding one bat that survived until the end of the experiment 50 days after the challenge). To do this, we calculated the mean behavior count for each focal rabid bat for a given period prior to death (e.g., 4 days = days 1-4 prior to death), then compared that observed mean to the expected mean (i.e., the mean count of all non-rabid bats within the same group and time period). To get an effect size for each time period, we standardized this mean difference:

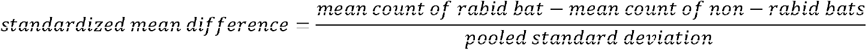

## Results

Fourteen bats died after the experimental RABV challenge; all were confirmed RABV positive, and deaths occurred in all three cages and in all three treatment groups (5 controls, 4 oral vaccinates, and 5 topically vaccinated). We detected no difference between the vaccination treatments on behavioral rates in rabid bats (Fig. S5, Table S6). The time of death ranged from 9-29 days post-challenge. One other RABV-challenged bat that had been topically vaccinated was alive by the end of the experiment (after 50 days) but confirmed rabid after it was euthanized. None of the 10 vaccinated bats that became rabid were shedding virus. In the unvaccinated rabid bats, we detected RABV shedding in the saliva of 3 bats on the day of death, and one of these was also shedding RABV the day prior to death.

In rabid bats, we did not detect a clear increase in aggression given (Fig. 1) or received (Fig. S2). Rabid vampire bats both gave and received less allogrooming than their healthy cagemates. This difference occurred on average about 12 days after inoculation and increased as we considered time periods closer to their death (Fig. 2, S3). The decrease in allogrooming and low levels of aggression are consistent with paralytic rather than furious rabies.

**Figure 1:**
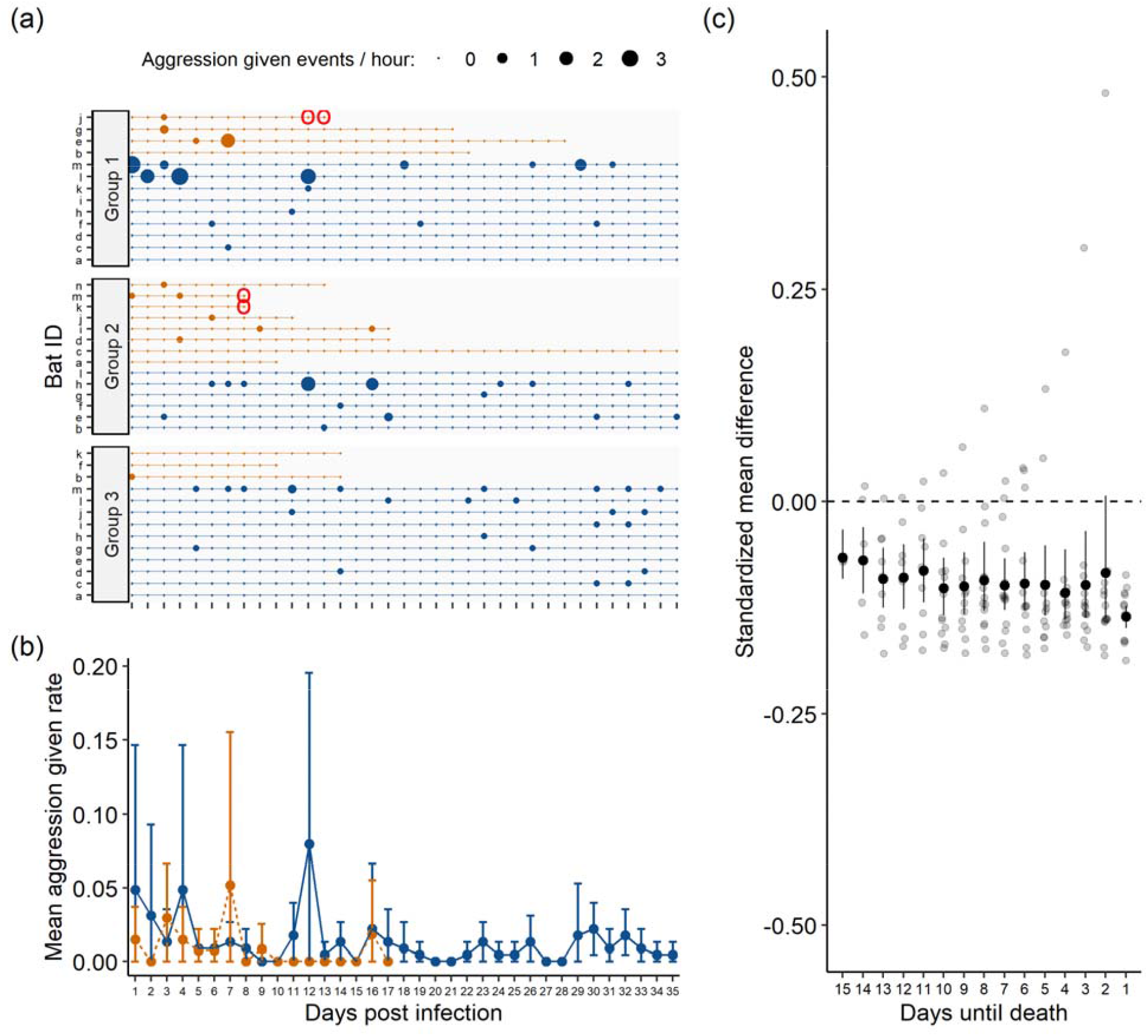
No evidence of increased aggression in rabid male vampire bats prior to death. Panel (a) shows timeline of aggression event counts for each rabid (orange points) and non-rabid (blue point) vampire bat across three groups. Point size reflects the rate of observed events per hour. Red circles show RABV positive saliva sample. Panel (b) shows mean rate of aggression events per hour with 95% CIs for rabid and non-rabid bats starting one day after inoculation with RABV. Panel (c) shows the standardized mean difference with 95% CIs between rabid bats and healthy cagemates during shrinking time intervals before death. Outliers in panel c are caused by one rabid bat (group 2-i) that showed aggression 16 days post-challenge (2 days prior to death). See Table S4 for CIs.

**Figure 2:**
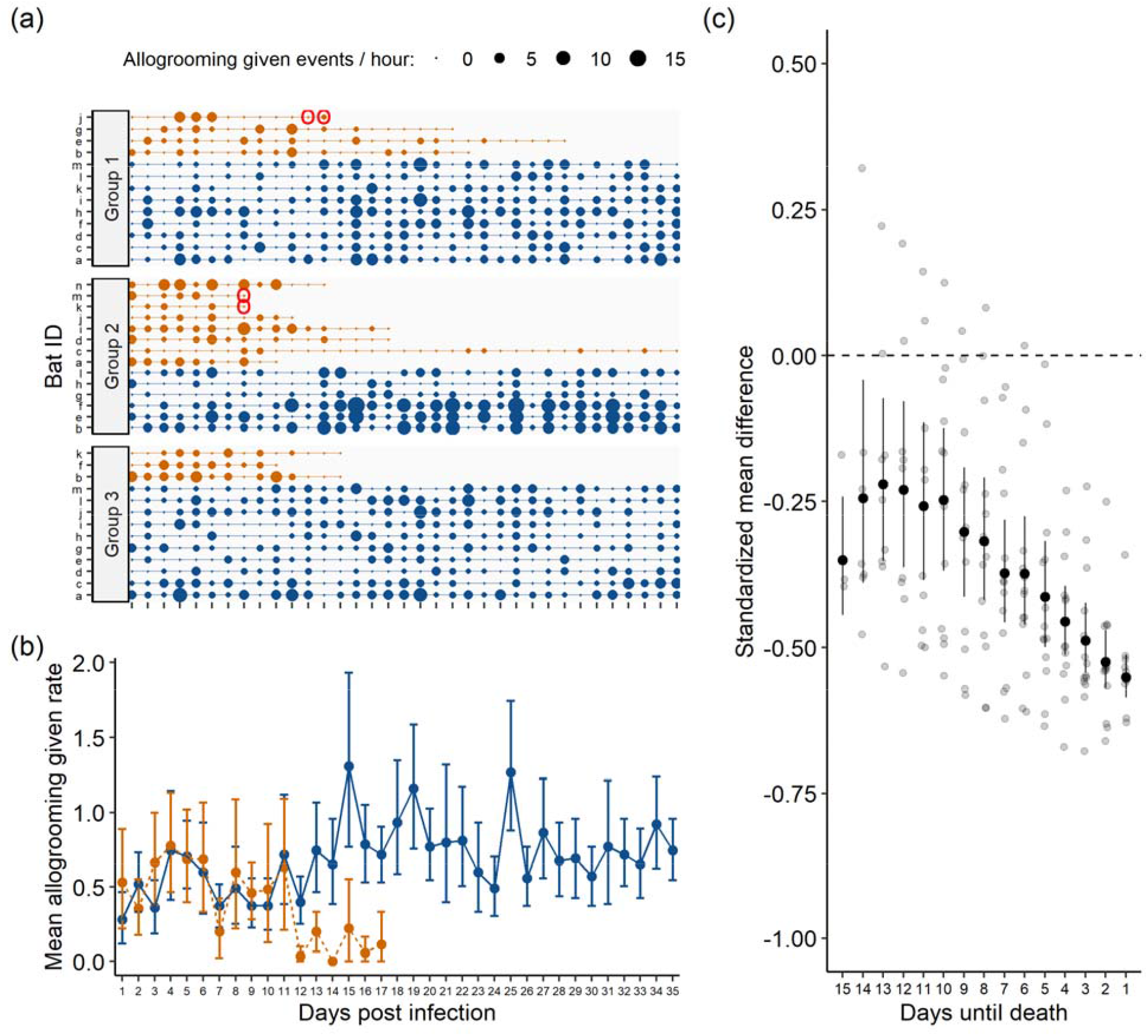
Reduced allogrooming prior to death in rabid male vampire bats. Panel (a) shows timeline of allogrooming events counts for each rabid (orange points) and non-rabid (blue point) vampire bat across three groups. Point size reflects the rate of observed events per hour. Red circles show RABV positive saliva sample. Panel (b) shows mean rate of allogrooming events per hour with 95% CIs for rabid and non-rabid bats starting one day after inoculation with RABV. Panel (c) shows the standardized mean difference with 95% CIs between rabid bats and healthy cagemates during shrinking time intervals before death. See Table S4 for CIs.

## Discussion

Rabid male vampire bats reduced their allogrooming, and probably as a consequence, also received less allogrooming. This change could be due to either rabies-induced paralysis or generalized sickness behaviors, such as lethargy, causing passive self-isolation [37–40]. All bats showed low rates of aggression, and we saw no clear increase in aggression in the rabid bats, regardless of vaccine treatment group.

Several other studies did not observe heightened aggression in rabid vampire bats (Table 1). One possible reason for this lack of observations is reduced selection on RABV to increase aggression in vampire bats because they are highly social, frequently aggressive, and bite other hosts [41]. Another possibility is that distinct RABV strains differ in pathogenicity and clinical forms of the disease (e.g., presence of aggression) across species [42–45]. Studies describing natural infections often report some aggression, but experimental RABV challenges that failed to find evidence of aggression used viral strains that were not currently circulating or, as in our case, used a strain derived from a different species (Table 1). It would be interesting to determine if infection with endemic vampire bat RABV strains may induce a higher proportion of furious versus paralytic disease in vampire bats.

Similar to field observations [17,36], some rabid bats in our study may have received increased aggression prior to death (see Fig. S2, e.g., bat group 1-j, 2-i, 2-d, 3-b). Acute infections can have a social cost, such as increased antagonism by conspecifics [46], and it would be interesting to examine evidence for avoidance of or aggression towards rabid individuals more closely in this and other species [40].

Times until death in rabid vampire bats varied from 9 to 29 days, but one of the 15 rabid males remained alive until the end of the experiment, 50 days after infection. The bat was previously vaccinated, but its neurologic function declined over the final weeks, losing coordination and mobility. We did not detect RABV in its saliva. The causes of this prolonged survival remain unclear.

In the late stage of infection, RABV spreads to the salivary glands and is excreted in saliva [42]. Evidence of RABV shedding in vampire bats prior to or at the time of death has been demonstrated before [20,33,47]. Here, we detected RABV shedding in saliva of 3 of 15 rabid bats (all 3 unvaccinated), which allowed us to overlay behavioral measures with pathogen shedding (Fig. 1, 2, S2, S3). These 3 vampire bats were not grooming others much when the virus was detectable in their saliva. Similarly, we did not observe heightened social aggression in these periods before death. Future work to quantify the relationship more closely between rabies shedding and behavioral changes would help distinguish this interaction.

Some caveats to the interpretations of our experimental results and those of others (Table 1) should be considered. First, given that social aggression can be rare and brief, absence of evidence of social aggression is not evidence of absence. We observed anecdotal evidence of aggression by some rabid bats towards handlers and other bats when the bats were disturbed. Second, the administered RABV challenge dose, route, and site of inoculation are not standardized across experiments and often differ (Table 1). As in our study, the RABV challenge strains used by researchers do not typically represent currently endemic viruses, are derived from different species, or are adapted to other species before their use in experimental infections. More standardized experimental infections are needed to disentangle the role of administered dose and temporal overlap of circulating strains on rabies-induced behavioral changes in natural reservoirs such as vampire bats.

In conclusion, we observed reductions in allogrooming and low levels of aggression that indicated paralytic but not furious rabies presentation in 15 rabid male vampire bats relative to 25 non-rabid male bats. Alongside other previous reports involving natural rabies exposures that report elevated aggression (Table 1), our results are consistent with the hypothesis that behavioral effects of RABV may vary by strain.

## Ethics

Field work was carried out under permit *SGPA/DGVS/003242/18* from the Mexican Secretariat of Environment and Natural Resources. All animal husbandry practices and experiments were approved by the USGS-National Wildlife Health Center Institutional Animal Care and Use Committee (*Protocol EP180418*). Any use of trade, firm, or product names is for descriptive purposes only and does not imply endorsement by the U.S. Government.

## Data Availability

All data and R code to repeat the analysis is publicly available on Figshare: https://doi.org/10.6084/m9.figshare.19991204.v3

## Acknowledgements

We thank Osric Wong, Natalie Sebunia, Jaelynn Butler, Michael Shechtman, Olivia Kalczynski, and Raven Hartman for video scoring. Daniel Streicker, Horacio Delpietro, Alvaro Aguilar-Setién, Charles Rupprecht provided literature. We thank Elizabeth Falendysz and Erik Hofmeister for manuscript review.

## Authors contributions

EMCC, SS, and GGC designed the study. EMCC carried out experiments. SS and EC analyzed the data. TR and JEO coordinated the study and provided resources and lab space. GGC conceived of the idea and supervised the data analysis and writing. All authors contributed to draft of manuscript and gave final approval for publication.

