## Supplemental materials for "Social effects of rabies infection in male vampire bats (*Desmodus rotundus*)"

**Supplementary materials for** “Social effects of rabies infection in male vampire bats (*Desmodus rotundus*)”

Sebastian Stockmaier*^2^

Eleanor Cronin^2^

Tonie Rocke^3^

Jorge E. Osorio^1^

Gerald G. Carter^2,4^

***co-first authors**

**Affiliations**

1. Department of Pathobiological Sciences, School of Veterinary Medicine, University of Wisconsin-Madison, Madison, WI 53706, USA
2. Department of Evolution, Ecology and Organismal Biology, The Ohio State University, Columbus, OH 43210, USA
3. U.S. Geological Survey, National Wildlife Health Center, Madison, WI 53711, USA
4. Smithsonian Tropical Research Institute, Balboa Ancón, Panama

**Disclaimer:** Any use of trade, firm, or product names is for descriptive purposes only and does not imply endorsement by the U.S. Government.

**Our manuscript includes three supplementary videos:**

**Video S1:** Video shows allogrooming in male vampire bats (*Desmodus rotundus*) which involves licking or chewing another bat’s fur or skin and often occurs with two bats allogrooming each other simultaneously. Video credit: U.S. Geological Survey National Wildlife Health Center/Department of Pathobiological Sciences, School of Veterinary Medicine, University of Wisconsin-Madison/Department of Evolution, Ecology, and Organismal Biology, The Ohio State University

**Video S2:** Video shows aggressive events including biting and fighting between two male vampire bats (*Desmodus rotundus*). Video credit: U.S. Geological Survey National Wildlife Health Center/Department of Pathobiological Sciences, School of Veterinary Medicine, University of Wisconsin-Madison/Department of Evolution, Ecology, and Organismal Biology, The Ohio State University

**Video S3:** Video shows clinging behavior where one male vampire bat (*Desmodus rotundus*) bites into another bats neck and clings onto it for a prolonged period while the target is actively trying to shake off the aggressor. Video credit: U.S. Geological Survey National Wildlife Health Center/Department of Pathobiological Sciences, School of Veterinary Medicine, University of Wisconsin-Madison/Department of Evolution, Ecology, and Organismal Biology, The Ohio State University

**Resumen S1**

El virus de la rabia (RABV) transmitido por el vampiro común (*Desmodus rotundus*) representa una amenaza para el desarrollo agrícola y la salud pública en todo el Neotrópico. La ecología y evolución de la dinámica hospedero-patógeno de la rabia están influenciadas por dos cambios de comportamiento inducidos por esta enfermedad. Frecuentemente, los hospederos con rabia exhiben una mayor agresión, lo cual facilita la transmisión del virus, aunque la rabia también puede inducir una actividad reducida y parálisis antes de la muerte. Aunque varios estudios han documentado cambios de comportamiento inducidos por la rabia en roedores y otros hospederos, sorprendentemente pocos estudios han medido estos cambios en el vampiro común, un importante reservorio natural de la rabia en toda Latinoamérica. En este estudio cuantificamos por primera vez cómo la rabia afecta los comportamientos de agresividad y acicalamiento en el murciélago vampiro, aprovechando el acceso a ejemplares machos en cautiverio, que formaban parte de un experimento de seguridad y eficacia de una vacuna oral contra la rabia. Los individuos con rabia redujeron su acicalamiento previo a la muerte en comparación con los vampiros no rabiosos, pero no detectamos aumentos en la agresión entre los murciélagos. Para poner nuestros resultados en contexto, hicimos una revisión de lo que es sabido y lo que aún queda por esclarecer sobre los cambios de comportamiento de los murciélagos vampiro con rabia.

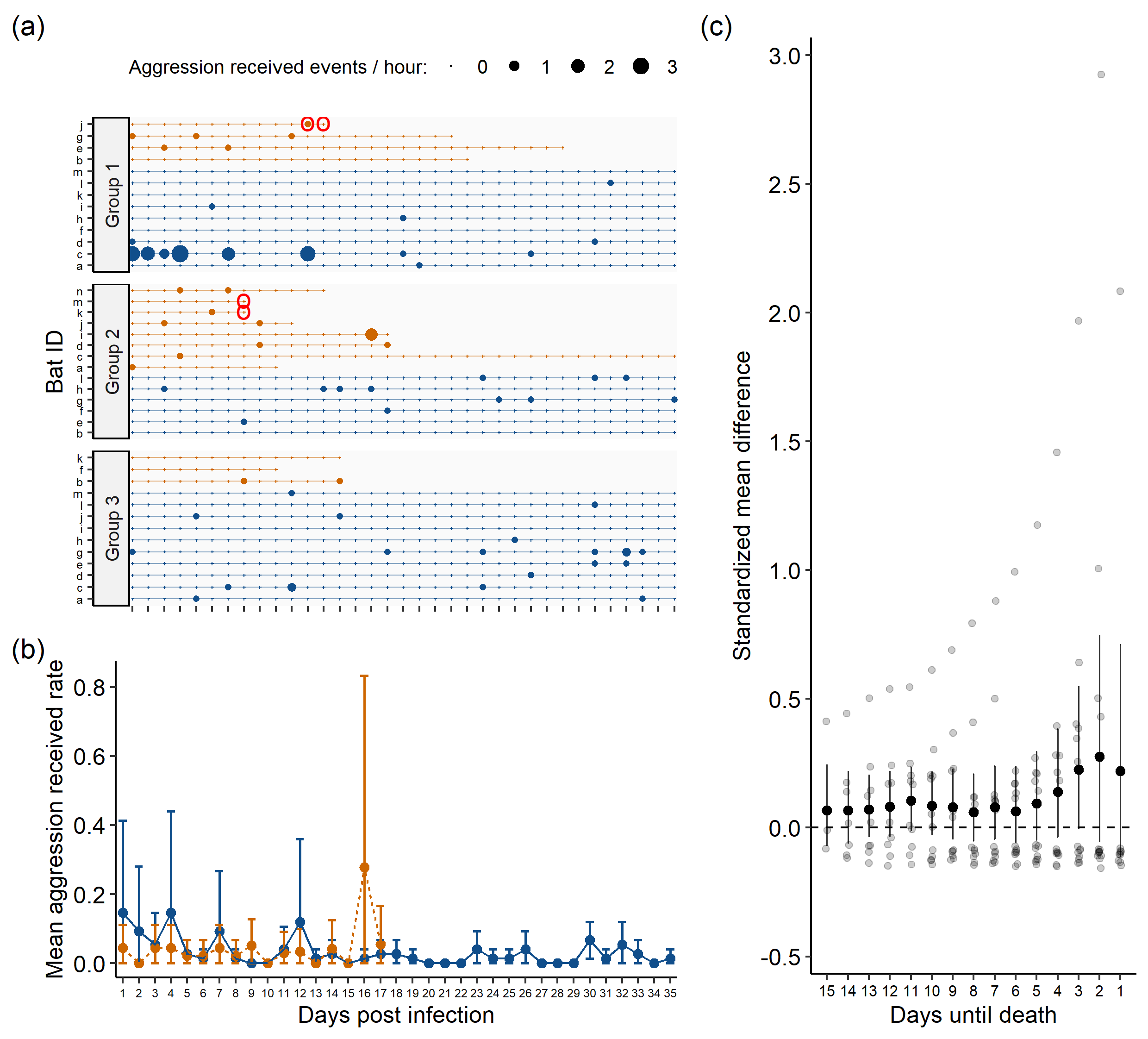

**Figure S2: No clear evidence of increased aggression towards rabid male vampire bats (*Desmodus rotundus*) prior to death.** Panel **(a)** shows timeline of aggression events towards a bat counted for each rabid (orange points) and non-rabid (blue point) vampire bat across three cages. Point size reflects the rate of observed events per hour and red circles show RABV positive saliva sample in bat. Panel **(b)** shows mean rate of aggression events per hour with 95% confidence intervals (CIs) towards rabid and non-rabid bats starting at one day after inoculation with RABV. Panel **(c)** shows the standardized mean difference with 95% CIs between rabid bats and healthy cagemates during shrinking time intervals before death. Outliers (>0) indicate bat that was targeted more by aggression prior to death than non-rabid bats during the same time period (e.g., bats group 1-j, 2-i, 2-d, 3-b). See Table S4 for CIs.

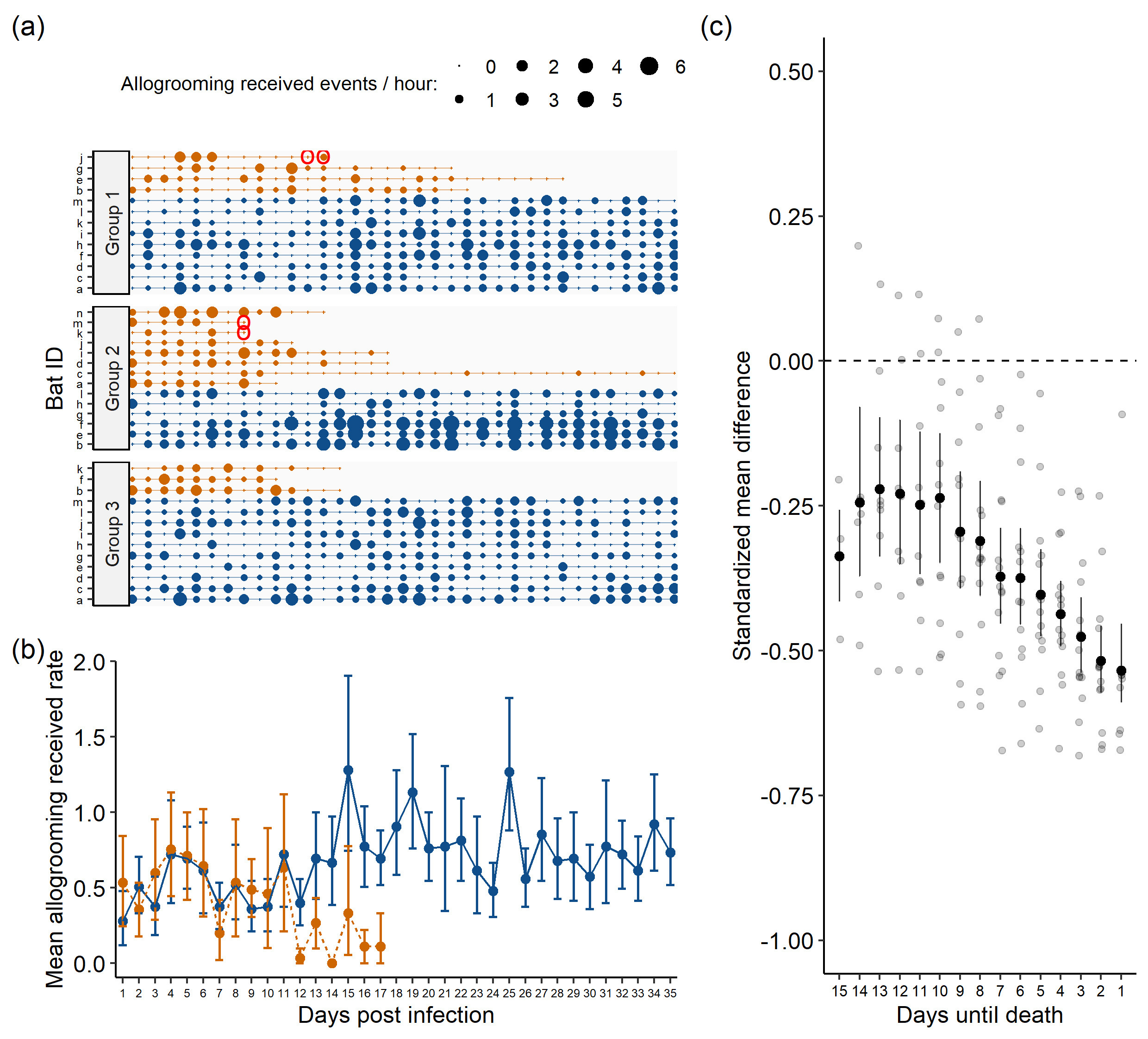

**Figure S3: Reduced allogrooming towards rabid male vampire bats (*Desmodus rotundus*) prior to death.** Panel **(a)** shows timeline of allogrooming events received by each rabid (orange points) and non-rabid (blue point) vampire bat across three cages. Point size reflects the rate of observed events per hour and red circles show RABV positive saliva sample in bat. Panel **(b)** shows mean rate of allogrooming events per hour with 95% CIs towards rabid and non-rabid bats starting at one day after inoculation with RABV. Panel **(c)** shows the standardized mean difference with 95% CIs between rabid bats and healthy cagemates during shrinking time intervals before death. See Table S4 for CIs.

**Table S4: Means and 95% bootstrapped confidence intervals (CI) for all presented figures.** We did not calculate confidence intervals for n < 5.

| Aggression towards others (Figure 1) | | | | | |
| --- | --- | --- | --- | --- | --- |
| *Days post infection* | *Rabid?* | *Lower CI* | *Mean* | *Upper CI* | *n* |
| 1 | FALSE | 0.00 | 0.05 | 0.15 | 25 |
| 1 | TRUE | 0.00 | 0.01 | 0.04 | 15 |
| 2 | FALSE | 0.00 | 0.03 | 0.09 | 25 |
| 2 | TRUE | 0.00 | 0.00 | 0.00 | 15 |
| 3 | FALSE | 0.00 | 0.01 | 0.04 | 25 |
| 3 | TRUE | 0.00 | 0.03 | 0.07 | 15 |
| 4 | FALSE | 0.00 | 0.05 | 0.15 | 25 |
| 4 | TRUE | 0.00 | 0.01 | 0.04 | 15 |
| 5 | FALSE | 0.00 | 0.01 | 0.02 | 25 |
| 5 | TRUE | 0.00 | 0.01 | 0.02 | 15 |
| 6 | FALSE | 0.00 | 0.01 | 0.02 | 25 |
| 6 | TRUE | 0.00 | 0.01 | 0.02 | 15 |
| 7 | FALSE | 0.00 | 0.01 | 0.03 | 25 |
| 7 | TRUE | 0.00 | 0.05 | 0.16 | 15 |
| 8 | FALSE | 0.00 | 0.01 | 0.02 | 25 |
| 8 | TRUE | 0.00 | 0.00 | 0.00 | 15 |
| 9 | FALSE | 0.00 | 0.00 | 0.00 | 25 |
| 9 | TRUE | 0.00 | 0.01 | 0.03 | 13 |
| 10 | FALSE | 0.00 | 0.00 | 0.00 | 25 |
| 10 | TRUE | 0.00 | 0.00 | 0.00 | 13 |
| 11 | FALSE | 0.00 | 0.02 | 0.04 | 25 |
| 11 | TRUE | 0.00 | 0.00 | 0.00 | 11 |
| 12 | FALSE | 0.00 | 0.08 | 0.20 | 25 |
| 12 | TRUE | 0.00 | 0.00 | 0.00 | 10 |
| 13 | FALSE | 0.00 | 0.00 | 0.01 | 25 |
| 13 | TRUE | 0.00 | 0.00 | 0.00 | 10 |
| 14 | FALSE | 0.00 | 0.01 | 0.03 | 25 |
| 14 | TRUE | 0.00 | 0.00 | 0.00 | 8 |
| 15 | FALSE | 0.00 | 0.00 | 0.00 | 25 |
| 15 | TRUE | 0.00 | 0.00 | 0.00 | 6 |
| 16 | FALSE | 0.00 | 0.02 | 0.07 | 25 |
| 16 | TRUE | 0.00 | 0.02 | 0.06 | 6 |
| 17 | FALSE | 0.00 | 0.01 | 0.04 | 25 |
| 17 | TRUE | 0.00 | 0.00 | 0.00 | 6 |
| 18 | FALSE | 0.00 | 0.01 | 0.03 | 25 |
| 19 | FALSE | 0.00 | 0.00 | 0.01 | 25 |
| 20 | FALSE | 0.00 | 0.00 | 0.00 | 25 |
| 21 | FALSE | 0.00 | 0.00 | 0.00 | 25 |
| 22 | FALSE | 0.00 | 0.00 | 0.01 | 25 |
| 23 | FALSE | 0.00 | 0.01 | 0.03 | 25 |
| 24 | FALSE | 0.00 | 0.00 | 0.01 | 25 |
| 25 | FALSE | 0.00 | 0.00 | 0.01 | 25 |
| 26 | FALSE | 0.00 | 0.01 | 0.03 | 25 |
| 27 | FALSE | 0.00 | 0.00 | 0.00 | 25 |
| 28 | FALSE | 0.00 | 0.00 | 0.00 | 25 |
| 29 | FALSE | 0.00 | 0.02 | 0.05 | 25 |
| 30 | FALSE | 0.01 | 0.02 | 0.04 | 25 |
| 31 | FALSE | 0.00 | 0.01 | 0.02 | 25 |
| 32 | FALSE | 0.00 | 0.02 | 0.04 | 25 |
| 33 | FALSE | 0.00 | 0.01 | 0.02 | 25 |
| 34 | FALSE | 0.00 | 0.00 | 0.01 | 25 |
| 35 | FALSE | 0.00 | 0.00 | 0.01 | 25 |
| Standardized mean difference (aggression towards others) | | | | | |
| *Time interval* |  | *Lower CI* | *Mean* | *Upper CI* | *n* |
| 1 |  | -0.15 | -0.14 | -0.12 | 14 |
| 2 |  | -0.14 | -0.08 | 0.01 | 14 |
| 3 |  | -0.14 | -0.10 | -0.03 | 14 |
| 4 |  | -0.14 | -0.11 | -0.06 | 14 |
| 5 |  | -0.13 | -0.10 | -0.05 | 14 |
| 6 |  | -0.13 | -0.10 | -0.06 | 14 |
| 7 |  | -0.13 | -0.10 | -0.07 | 14 |
| 8 |  | -0.13 | -0.09 | -0.05 | 14 |
| 9 |  | -0.13 | -0.10 | -0.06 | 12 |
| 10 |  | -0.13 | -0.10 | -0.07 | 12 |
| 11 |  | -0.12 | -0.08 | -0.04 | 10 |
| 12 |  | -0.13 | -0.09 | -0.05 | 9 |
| 13 |  | -0.13 | -0.09 | -0.06 | 9 |
| 14 |  | -0.11 | -0.07 | -0.03 | 7 |
| 15 |  | -0.09 | -0.07 | -0.03 | 5 |
| Allogrooming given (Figure 2) | | | | | |
| *Days post infection* | *Rabid?* | *Lower CI* | *Mean* | *Upper CI* | *n* |
| 1 | FALSE | 0.12 | 0.28 | 0.47 | 25 |
| 1 | TRUE | 0.22 | 0.53 | 0.89 | 15 |
| 2 | FALSE | 0.33 | 0.52 | 0.72 | 25 |
| 2 | TRUE | 0.18 | 0.36 | 0.56 | 15 |
| 3 | FALSE | 0.19 | 0.36 | 0.56 | 25 |
| 3 | TRUE | 0.38 | 0.67 | 1.02 | 15 |
| 4 | FALSE | 0.41 | 0.75 | 1.15 | 25 |
| 4 | TRUE | 0.47 | 0.78 | 1.13 | 15 |
| 5 | FALSE | 0.51 | 0.71 | 0.93 | 25 |
| 5 | TRUE | 0.40 | 0.69 | 1.00 | 15 |
| 6 | FALSE | 0.32 | 0.60 | 0.92 | 25 |
| 6 | TRUE | 0.36 | 0.69 | 1.07 | 15 |
| 7 | FALSE | 0.23 | 0.37 | 0.53 | 25 |
| 7 | TRUE | 0.02 | 0.20 | 0.42 | 15 |
| 8 | FALSE | 0.27 | 0.49 | 0.77 | 25 |
| 8 | TRUE | 0.22 | 0.60 | 1.09 | 15 |
| 9 | FALSE | 0.23 | 0.37 | 0.55 | 25 |
| 9 | TRUE | 0.28 | 0.46 | 0.67 | 13 |
| 10 | FALSE | 0.21 | 0.37 | 0.56 | 25 |
| 10 | TRUE | 0.13 | 0.49 | 0.95 | 13 |
| 11 | FALSE | 0.39 | 0.72 | 1.13 | 25 |
| 11 | TRUE | 0.21 | 0.64 | 1.09 | 11 |
| 12 | FALSE | 0.25 | 0.40 | 0.56 | 25 |
| 12 | TRUE | 0.00 | 0.03 | 0.10 | 10 |
| 13 | FALSE | 0.47 | 0.75 | 1.07 | 25 |
| 13 | TRUE | 0.07 | 0.20 | 0.33 | 10 |
| 14 | FALSE | 0.39 | 0.65 | 0.96 | 25 |
| 14 | TRUE | 0.00 | 0.00 | 0.00 | 8 |
| 15 | FALSE | 0.77 | 1.31 | 1.93 | 25 |
| 15 | TRUE | 0.00 | 0.22 | 0.56 | 6 |
| 16 | FALSE | 0.53 | 0.79 | 1.05 | 25 |
| 16 | TRUE | 0.00 | 0.06 | 0.17 | 6 |
| 17 | FALSE | 0.53 | 0.72 | 0.92 | 25 |
| 17 | TRUE | 0.00 | 0.11 | 0.33 | 6 |
| 18 | FALSE | 0.60 | 0.93 | 1.35 | 25 |
| 19 | FALSE | 0.76 | 1.16 | 1.60 | 25 |
| 20 | FALSE | 0.55 | 0.77 | 1.03 | 25 |
| 21 | FALSE | 0.39 | 0.80 | 1.31 | 25 |
| 22 | FALSE | 0.51 | 0.81 | 1.15 | 25 |
| 23 | FALSE | 0.32 | 0.60 | 0.92 | 25 |
| 24 | FALSE | 0.31 | 0.49 | 0.72 | 25 |
| 25 | FALSE | 0.87 | 1.27 | 1.76 | 25 |
| 26 | FALSE | 0.36 | 0.56 | 0.76 | 25 |
| 27 | FALSE | 0.56 | 0.87 | 1.24 | 25 |
| 28 | FALSE | 0.44 | 0.68 | 0.93 | 25 |
| 29 | FALSE | 0.43 | 0.69 | 0.97 | 25 |
| 30 | FALSE | 0.37 | 0.57 | 0.79 | 25 |
| 31 | FALSE | 0.39 | 0.77 | 1.23 | 25 |
| 32 | FALSE | 0.49 | 0.72 | 0.96 | 25 |
| 33 | FALSE | 0.43 | 0.65 | 0.89 | 25 |
| 34 | FALSE | 0.63 | 0.92 | 1.24 | 25 |
| 35 | FALSE | 0.55 | 0.75 | 0.96 | 25 |
| Standardized mean difference (allogrooming given) | | | | | |
| *Time interval* |  | *Lower CI* | *Mean* | *Upper CI* | *n* |
| 1 |  | -0.58 | -0.55 | -0.51 | 14 |
| 2 |  | -0.57 | -0.52 | -0.47 | 14 |
| 3 |  | -0.55 | -0.49 | -0.42 | 14 |
| 4 |  | -0.51 | -0.46 | -0.40 | 14 |
| 5 |  | -0.50 | -0.41 | -0.32 | 14 |
| 6 |  | -0.46 | -0.37 | -0.27 | 14 |
| 7 |  | -0.46 | -0.37 | -0.28 | 14 |
| 8 |  | -0.41 | -0.32 | -0.21 | 14 |
| 9 |  | -0.41 | -0.30 | -0.19 | 12 |
| 10 |  | -0.37 | -0.25 | -0.13 | 12 |
| 11 |  | -0.39 | -0.26 | -0.12 | 10 |
| 12 |  | -0.37 | -0.23 | -0.08 | 9 |
| 13 |  | -0.35 | -0.22 | -0.08 | 9 |
| 14 |  | -0.39 | -0.24 | -0.05 | 7 |
| 15 |  | -0.44 | -0.35 | -0.24 | 5 |
| Aggression received (Figure S2) | | | | | |
| *Days post infection* | *Rabid?* | *Lower CI* | *Mean* | *Upper CI* | *n* |
| 1 | FALSE | 0.00 | 0.15 | 0.40 | 25 |
| 1 | TRUE | 0.00 | 0.04 | 0.11 | 15 |
| 2 | FALSE | 0.00 | 0.09 | 0.28 | 25 |
| 2 | TRUE | 0.00 | 0.00 | 0.00 | 15 |
| 3 | FALSE | 0.00 | 0.05 | 0.15 | 25 |
| 3 | TRUE | 0.00 | 0.04 | 0.11 | 15 |
| 4 | FALSE | 0.00 | 0.15 | 0.44 | 25 |
| 4 | TRUE | 0.00 | 0.04 | 0.11 | 15 |
| 5 | FALSE | 0.00 | 0.03 | 0.07 | 25 |
| 5 | TRUE | 0.00 | 0.02 | 0.07 | 15 |
| 6 | FALSE | 0.00 | 0.01 | 0.04 | 25 |
| 6 | TRUE | 0.00 | 0.02 | 0.07 | 15 |
| 7 | FALSE | 0.00 | 0.09 | 0.27 | 25 |
| 7 | TRUE | 0.00 | 0.04 | 0.11 | 15 |
| 8 | FALSE | 0.00 | 0.01 | 0.04 | 25 |
| 8 | TRUE | 0.00 | 0.02 | 0.07 | 15 |
| 9 | FALSE | 0.00 | 0.00 | 0.00 | 25 |
| 9 | TRUE | 0.00 | 0.05 | 0.13 | 13 |
| 10 | FALSE | 0.00 | 0.00 | 0.00 | 25 |
| 10 | TRUE | 0.00 | 0.00 | 0.00 | 13 |
| 11 | FALSE | 0.00 | 0.04 | 0.11 | 25 |
| 11 | TRUE | 0.00 | 0.03 | 0.09 | 11 |
| 12 | FALSE | 0.00 | 0.12 | 0.36 | 25 |
| 12 | TRUE | 0.00 | 0.03 | 0.10 | 10 |
| 13 | FALSE | 0.00 | 0.01 | 0.04 | 25 |
| 13 | TRUE | 0.00 | 0.00 | 0.00 | 10 |
| 14 | FALSE | 0.00 | 0.03 | 0.07 | 25 |
| 14 | TRUE | 0.00 | 0.04 | 0.12 | 8 |
| 15 | FALSE | 0.00 | 0.00 | 0.00 | 25 |
| 15 | TRUE | 0.00 | 0.00 | 0.00 | 6 |
| 16 | FALSE | 0.00 | 0.01 | 0.04 | 25 |
| 16 | TRUE | 0.00 | 0.28 | 0.83 | 6 |
| 17 | FALSE | 0.00 | 0.03 | 0.07 | 25 |
| 17 | TRUE | 0.00 | 0.06 | 0.17 | 6 |
| 18 | FALSE | 0.00 | 0.03 | 0.07 | 25 |
| 19 | FALSE | 0.00 | 0.01 | 0.04 | 25 |
| 20 | FALSE | 0.00 | 0.00 | 0.00 | 25 |
| 21 | FALSE | 0.00 | 0.00 | 0.00 | 25 |
| 22 | FALSE | 0.00 | 0.00 | 0.00 | 25 |
| 23 | FALSE | 0.00 | 0.04 | 0.08 | 25 |
| 24 | FALSE | 0.00 | 0.01 | 0.04 | 25 |
| 25 | FALSE | 0.00 | 0.01 | 0.04 | 25 |
| 26 | FALSE | 0.00 | 0.04 | 0.09 | 25 |
| 27 | FALSE | 0.00 | 0.00 | 0.00 | 25 |
| 28 | FALSE | 0.00 | 0.00 | 0.00 | 25 |
| 29 | FALSE | 0.00 | 0.00 | 0.00 | 25 |
| 30 | FALSE | 0.01 | 0.07 | 0.12 | 25 |
| 31 | FALSE | 0.00 | 0.01 | 0.04 | 25 |
| 32 | FALSE | 0.00 | 0.05 | 0.12 | 25 |
| 33 | FALSE | 0.00 | 0.03 | 0.07 | 25 |
| 34 | FALSE | 0.00 | 0.00 | 0.00 | 25 |
| 35 | FALSE | 0.00 | 0.01 | 0.04 | 25 |
| Standardized mean difference (aggression received) | | | | | |
| *Time interval* |  | *Lower CI* | *Mean* | *Upper CI* | *n* |
| 1 |  | -0.11 | 0.22 | 0.71 | 14 |
| 2 |  | -0.06 | 0.28 | 0.75 | 14 |
| 3 |  | -0.01 | 0.23 | 0.55 | 14 |
| 4 |  | -0.03 | 0.14 | 0.39 | 14 |
| 5 |  | -0.05 | 0.09 | 0.29 | 14 |
| 6 |  | -0.06 | 0.06 | 0.23 | 14 |
| 7 |  | -0.05 | 0.08 | 0.24 | 14 |
| 8 |  | -0.05 | 0.06 | 0.20 | 14 |
| 9 |  | -0.05 | 0.08 | 0.23 | 12 |
| 10 |  | -0.03 | 0.09 | 0.22 | 12 |
| 11 |  | -0.01 | 0.11 | 0.24 | 10 |
| 12 |  | -0.04 | 0.08 | 0.22 | 9 |
| 13 |  | -0.04 | 0.07 | 0.21 | 9 |
| 14 |  | -0.06 | 0.07 | 0.22 | 7 |
| 15 |  | -0.07 | 0.07 | 0.25 | 5 |
| Allogrooming received (Figure S3) | | | | | |
| *Days post infection* | *Rabid?* | *Lower CI* | *Mean* | *Upper CI* | *n* |
| 1 | FALSE | 0.12 | 0.28 | 0.47 | 25 |
| 1 | TRUE | 0.25 | 0.53 | 0.84 | 15 |
| 2 | FALSE | 0.33 | 0.51 | 0.69 | 25 |
| 2 | TRUE | 0.18 | 0.36 | 0.53 | 15 |
| 3 | FALSE | 0.20 | 0.37 | 0.57 | 25 |
| 3 | TRUE | 0.31 | 0.60 | 0.93 | 15 |
| 4 | FALSE | 0.40 | 0.72 | 1.09 | 25 |
| 4 | TRUE | 0.44 | 0.76 | 1.13 | 15 |
| 5 | FALSE | 0.49 | 0.69 | 0.92 | 25 |
| 5 | TRUE | 0.42 | 0.71 | 1.00 | 15 |
| 6 | FALSE | 0.32 | 0.61 | 0.92 | 25 |
| 6 | TRUE | 0.33 | 0.64 | 1.00 | 15 |
| 7 | FALSE | 0.23 | 0.37 | 0.53 | 25 |
| 7 | TRUE | 0.02 | 0.20 | 0.42 | 15 |
| 8 | FALSE | 0.28 | 0.52 | 0.79 | 25 |
| 8 | TRUE | 0.18 | 0.53 | 0.93 | 15 |
| 9 | FALSE | 0.21 | 0.36 | 0.55 | 25 |
| 9 | TRUE | 0.31 | 0.49 | 0.69 | 13 |
| 10 | FALSE | 0.21 | 0.37 | 0.56 | 25 |
| 10 | TRUE | 0.13 | 0.46 | 0.87 | 13 |
| 11 | FALSE | 0.40 | 0.72 | 1.13 | 25 |
| 11 | TRUE | 0.21 | 0.64 | 1.12 | 11 |
| 12 | FALSE | 0.25 | 0.40 | 0.56 | 25 |
| 12 | TRUE | 0.00 | 0.03 | 0.10 | 10 |
| 13 | FALSE | 0.44 | 0.69 | 1.01 | 25 |
| 13 | TRUE | 0.13 | 0.27 | 0.43 | 10 |
| 14 | FALSE | 0.40 | 0.67 | 0.97 | 25 |
| 14 | TRUE | 0.00 | 0.00 | 0.00 | 8 |
| 15 | FALSE | 0.76 | 1.28 | 1.91 | 25 |
| 15 | TRUE | 0.06 | 0.33 | 0.78 | 6 |
| 16 | FALSE | 0.53 | 0.77 | 1.04 | 25 |
| 16 | TRUE | 0.00 | 0.11 | 0.22 | 6 |
| 17 | FALSE | 0.52 | 0.69 | 0.88 | 25 |
| 17 | TRUE | 0.00 | 0.11 | 0.33 | 6 |
| 18 | FALSE | 0.60 | 0.91 | 1.28 | 25 |
| 19 | FALSE | 0.76 | 1.13 | 1.53 | 25 |
| 20 | FALSE | 0.55 | 0.76 | 1.00 | 25 |
| 21 | FALSE | 0.35 | 0.77 | 1.31 | 25 |
| 22 | FALSE | 0.53 | 0.81 | 1.11 | 25 |
| 23 | FALSE | 0.32 | 0.61 | 0.97 | 25 |
| 24 | FALSE | 0.31 | 0.48 | 0.67 | 25 |
| 25 | FALSE | 0.89 | 1.27 | 1.72 | 25 |
| 26 | FALSE | 0.39 | 0.56 | 0.76 | 25 |
| 27 | FALSE | 0.56 | 0.85 | 1.23 | 25 |
| 28 | FALSE | 0.43 | 0.68 | 0.96 | 25 |
| 29 | FALSE | 0.41 | 0.69 | 1.00 | 25 |
| 30 | FALSE | 0.36 | 0.57 | 0.80 | 25 |
| 31 | FALSE | 0.39 | 0.77 | 1.24 | 25 |
| 32 | FALSE | 0.49 | 0.72 | 0.95 | 25 |
| 33 | FALSE | 0.40 | 0.61 | 0.84 | 25 |
| 34 | FALSE | 0.60 | 0.92 | 1.24 | 25 |
| 35 | FALSE | 0.52 | 0.73 | 0.96 | 25 |
| Standardized mean difference (allogrooming received) | | | | | |
| *Time interval* |  | *Lower CI* | *Mean* | *Upper CI* | *n* |
| 1 |  | -0.59 | -0.53 | -0.45 | 14 |
| 2 |  | -0.57 | -0.52 | -0.45 | 14 |
| 3 |  | -0.54 | -0.48 | -0.41 | 14 |
| 4 |  | -0.49 | -0.44 | -0.38 | 14 |
| 5 |  | -0.48 | -0.40 | -0.32 | 14 |
| 6 |  | -0.46 | -0.37 | -0.29 | 14 |
| 7 |  | -0.45 | -0.37 | -0.29 | 14 |
| 8 |  | -0.40 | -0.31 | -0.22 | 14 |
| 9 |  | -0.40 | -0.29 | -0.19 | 12 |
| 10 |  | -0.35 | -0.24 | -0.12 | 12 |
| 11 |  | -0.37 | -0.25 | -0.12 | 10 |
| 12 |  | -0.35 | -0.23 | -0.10 | 9 |
| 13 |  | -0.34 | -0.22 | -0.10 | 9 |
| 14 |  | -0.37 | -0.24 | -0.07 | 7 |
| 15 |  | -0.43 | -0.34 | -0.26 | 5 |

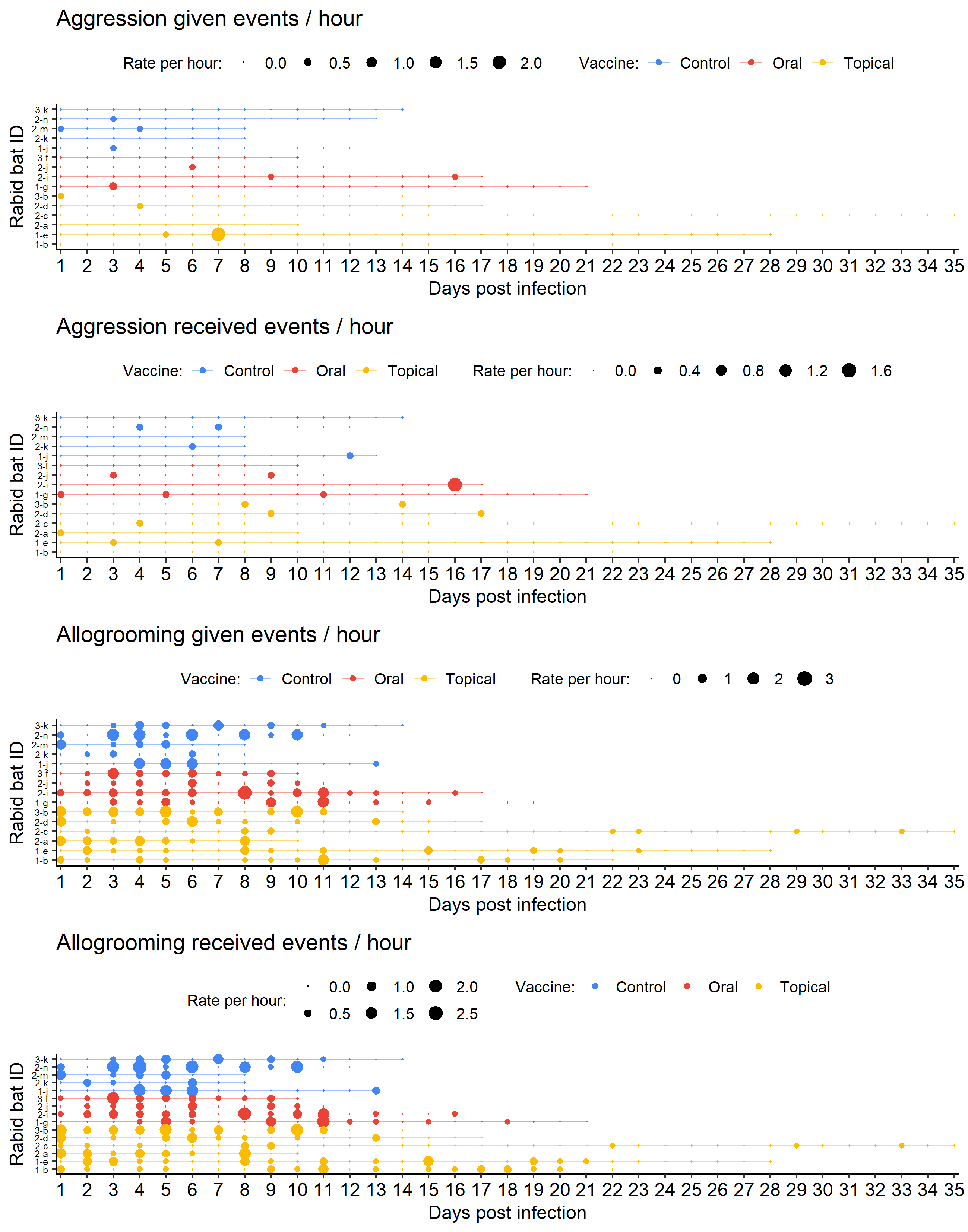

**Figure S5: No clear effects of previous vaccination status on behavioral rates in rabid vampire bats (*Desmodus rotundus*).** Bats were vaccinated with three different vaccine types (oral, topical, control) before rabies challenge and behavioral observations (Rabid bat id = “group – id”). Plots show behavioral rates per hour of all rabid bats based on previous vaccination treatment.

**Table S6: Model summary.** Vampire bats were vaccinated with three different vaccine types (oral, topical, placebo control) before rabies challenge and behavioral observations. We tested for effects of previous vaccination on behavioral rates per hour (aggression given, aggression received, allogrooming, allogrooming received) using linear mixed effect models. Rates were response variables and we tested for interaction effects of days post challenge and vaccination treatment (control, oral, topical). Bat ID was a random intercept. Overall, we did not detect an effect of previous vaccination on behavioral rates during the experiment.

|  | **Aggression given** | | **Aggression received** | | **Allogrooming given** | | **Allogrooming received** | |
| --- | --- | --- | --- | --- | --- | --- | --- | --- |
| *Predictors* | *Estimates*  *(95% CI)* | *P-Value* | *Estimates*  *(95% CI)* | *P-Value* | *Estimates*  *(95% CI)* | *P-Value* | *Estimates*  *(95% CI)* | *P-Value* |
| Intercept | 0.06 (-0.02 – 0.15) | 0.156 | 0.01 (-0.06 – 0.09) | 0.694 | 0.76 (0.41 – 1.10) | **<0.001** | 0.74 (0.43 – 1.05) | **<0.001** |
| Days post challenge | -0.01 (-0.02 – 0.01) | 0.298 | 0.00 (-0.01 – 0.01) | 0.765 | -0.05 (-0.08 – -0.01) | **0.024** | -0.04 (-0.08 – -0.00) | **0.037** |
| Oral vaccine | -0.03 (-0.15 – 0.10) | 0.668 | 0.00 (-0.10 – 0.10) | 0.968 | -0.08 (-0.56 – 0.40) | 0.749 | -0.11 (-0.55 – 0.33) | 0.630 |
| Topical vaccine | -0.01 (-0.11 – 0.10) | 0.893 | 0.02 (-0.06 – 0.11) | 0.563 | -0.17 (-0.59 – 0.25) | 0.415 | -0.17 (-0.55 – 0.21) | 0.388 |
| Interaction: Days post challenge x Oral vaccine | 0.01 (-0.01 – 0.02) | 0.483 | 0.00 (-0.01 – 0.01) | 0.589 | 0.02 (-0.03 – 0.07) | 0.441 | 0.02 (-0.03 – 0.07) | 0.427 |
| Interaction: Days post challenge x Topical vaccine | 0.00 (-0.01 – 0.02) | 0.540 | -0.00 (-0.01 – 0.01) | 0.576 | 0.03 (-0.01 – 0.07) | 0.169 | 0.02 (-0.02 – 0.06) | 0.242 |
| **Random Effects** | | | | | | | | |
| σ^2^ | 0.03 | | 0.02 | | 0.28 | | 0.27 | |
| τ_00_ | 0.00 _id_ | | 0.00 _id_ | | 0.05 _id_ | | 0.03 _id_ | |
| N | 15 _id_ | | 15 _id_ | | 15 _id_ | | 15 _id_ | |
| Observations | 241 | | 241 | | 241 | | 241 | |
| Marginal R^2^ / Conditional R^2^ | 0.013 / 0.016 | | 0.024 / NA | | 0.068 / 0.199 | | 0.070 / 0.157 | |
